# Ultrasonic Neuromodulation of the Human Hippocampus Reveals a Causal Role in Rapid Motor Memory Consolidation

**DOI:** 10.64898/2025.12.10.693529

**Authors:** Maria Paz Montenegro, Shancheng Bao, Sapna Bisht, Kenneth L. Hoyt, Huiliang Wang, David L. Wright, John J. Buchanan, Yuming Lei

## Abstract

How humans acquire motor skills is a fundamental biological question. Early learning involves rapid gains during practice and brief rest, yet the causal contribution of the hippocampus remains unclear. Here, we applied individualized, MRI-guided transcranial ultrasound stimulation (TUS) to modulate the posterior hippocampus in healthy adults. To establish a physiological benchmark, we first showed that this TUS protocol produced a sustained reduction in corticospinal excitability when applied to the primary motor cortex. When applied to the posterior hippocampus, electroencephalography revealed reduced frontocentral gamma-band activity, providing physiological evidence of hippocampal–frontal network modulation. Behaviorally, TUS altered the temporal dynamics of skill acquisition. Compared with sham, hippocampal TUS attenuated micro-offline gains during rest while increasing performance improvements during active practice. This shift from offline to online learning was replicated in an independent cohort with comparable baseline motor performance. Importantly, hippocampal TUS enhanced overall performance during learning, and this higher level of performance remained evident 24 hours later. Together, these findings provide causal evidence that the human hippocampus contributes to how motor learning is distributed between active practice and brief rest, showing that this temporal organization of learning can be altered by non-invasive neuromodulation.

## Introduction

The ability to learn motor skills through practice is fundamental to human achievement, from mastering daily routines to developing expert levels of performance (Dayan and Cohen, 2011; Krakauer et al., 2019). Central to this process is offline memory consolidation, during which fragile new memories are stabilized into more durable forms that are resistant to decay and interference (Walker et al., 2002; Robertson et al., 2004; Dudai, 2004; Frankland and Bontempi, 2005; Bao et al., 2022; Wang et al., 2024). In addition to the well-established process of memory consolidation that unfolds over hours and often involves sleep (Walker et al., 2002; Dudai and Eisenberg, 2004; Diekelmann and Born, 2010; Rasch and Born, 2013), recent studies have identified a rapid form of consolidation operating on a timescale of seconds. This “micro-consolidation,” occurring during brief rests interspersed with practice, appears to contribute substantially to the initial stages of learning (Bönstrup et al., 2019; Jacobacci et al., 2020; Buch et al., 2021).

Several lines of evidence suggest that the hippocampus is an important neural substrate for this rapid micro-consolidation. Neuroimaging studies show that heightened hippocampal activity and hippocampo-neocortical replay during these brief rests predict subsequent micro-offline performance gains (Jacobacci et al., 2020; Buch et al., 2021), while the hippocampus also exhibits rapid structural and functional plasticity during the learning of motor sequences (Jacobacci et al., 2020). Moreover, studies involving prefrontal stimulation that alters hippocampal reactivation (Gann et al., 2023) and patients with hippocampal damage (Mylonas et al., 2024) have shown attenuated micro-offline benefits. Together, these findings support a role for the hippocampus, but the evidence remains indirect. Neuroimaging provides correlational rather than causal evidence, while lesion and network disruption studies cannot isolate the hippocampus from the confounding effects of pathology or compensatory mechanisms. Thus, establishing a causal link requires an approach that allows non-invasive and targeted modulation of deep hippocampal regions in humans.

Directly testing this question remains difficult with conventional neuromodulation technologies. Non-invasive techniques such as transcranial magnetic stimulation (TMS) and transcranial electrical stimulation (tES) are limited in spatial focality and penetration depth (Polanía et al., 2018). These limitations make it difficult to selectively engage deep targets like the hippocampus without causing off-target stimulation in the overlying cortex, thereby confounding causal interpretation (Thielscher and Kammer, 2004; Bikson et al., 2016). Invasive methods such as deep brain stimulation (DBS) offer the necessary precision to target key structures within hippocampal circuits (Hamani et al., 2008; Suthana et al., 2012; Lozano and Lipsman, 2013), but their use in healthy populations is not feasible because of significant safety risks, including hemorrhage and infection (Hariz, 2002; Fenoy and Simpson, 2014). As a result, the causal role of deep brain structures in human motor learning has remained largely inaccessible to direct investigation.

Transcranial ultrasound stimulation (TUS) offers a way to overcome this methodological gap. TUS uses focused acoustic energy to non-invasively reach deep brain regions with greater spatial focality than conventional non-invasive stimulation approaches (Tyler et al., 2008; Legon et al., 2014; Fomenko et al., 2018; Blackmore et al., 2019). Its neuromodulatory effects likely arise from controlled mechanical interactions between acoustic waves and neural tissue, which may engage mechanosensitive ion channels to produce both transient and long-lasting changes in local neural activity (Yoo et al., 2011; Zeng et al., 2022; Song et al., 2023; Bao et al., 2024). Studies across preclinical and clinical settings have shown that TUS can modulate neural activity in targeted brain regions (Tufail et al., 2010; Monti et al., 2016; Legon et al., 2018; Folloni et al., 2019; Pasquinelli et al., 2019; Sanguinetti et al., 2020; Fomenko et al., 2020; Hou and Lei, 2025). Recent consensus guidelines also support its safety when stimulation parameters remain within established biophysical limits (Aubry et al., 2025). Ultrasonic neuromodulation has also been shown to influence learning- and memory-related processes in both animals (Eguchi et al., 2018) and humans (Jung et al., 2026). These properties make TUS particularly suitable for testing the causal contribution of the hippocampus to rapid motor memory consolidation in humans.

Here, we applied low-intensity TUS to directly test the causal role of the human hippocampus in rapid motor memory consolidation. We targeted its posterior segment, given its proposed role in spatial cognition and the temporal organization of sequential actions (Poppenk et al., 2013; Strange et al., 2014). Our approach involved several validation and experimental steps. First, we conducted acoustic validation with hydrophone measurements to verify our computational models, which were then used for individualized TUS targeting based on acoustic simulations and neuroanatomical imaging. Second, because the deep location of the hippocampus makes direct, non-invasive measurement of local cellular activity difficult, we established a quantifiable physiological benchmark for the neuromodulatory effect of our stimulation protocol. By applying the same TUS parameters to the primary motor cortex (M1), we showed that the protocol produced sustained suppression of corticospinal excitability, as assessed by motor-evoked potentials (MEPs). Third, having validated our neuromodulatory protocol, we used electroencephalography (EEG) to examine whether targeted hippocampal TUS altered cortical activity. Hippocampal TUS reduced frontocentral gamma-band activity, providing physiological evidence of hippocampal–frontal network modulation. Finally, we examined the behavioral impact of personalized hippocampal TUS on motor sequence learning in healthy participants using a motor sequence-learning protocol adapted from Bönstrup et al. (2019). We found that hippocampal TUS attenuated typical micro-offline gains (offline learning) while shifting performance improvements toward active practice (online learning). This shift in learning dynamics was reproduced in an independent cohort in which baseline motor performance was comparable between groups. Importantly, hippocampal TUS also enhanced overall task performance during initial learning, and this higher performance remained evident 24 hours later.

## Results

To evaluate the neurophysiological and behavioral impact of hippocampal TUS, participants underwent a randomized between-subjects design, receiving either active or sham stimulation prior to a motor sequence learning task. As depicted in Figure 1A, the primary behavioral paradigm consisted of 36 learning trials on Day 1, followed by a 9-trial retention test approximately 24 hours later on Day 2. Each trial comprised a 10-second active practice window, during which participants typed a five-element sequence, interleaved with a 10-second rest period.

**Figure 1.**
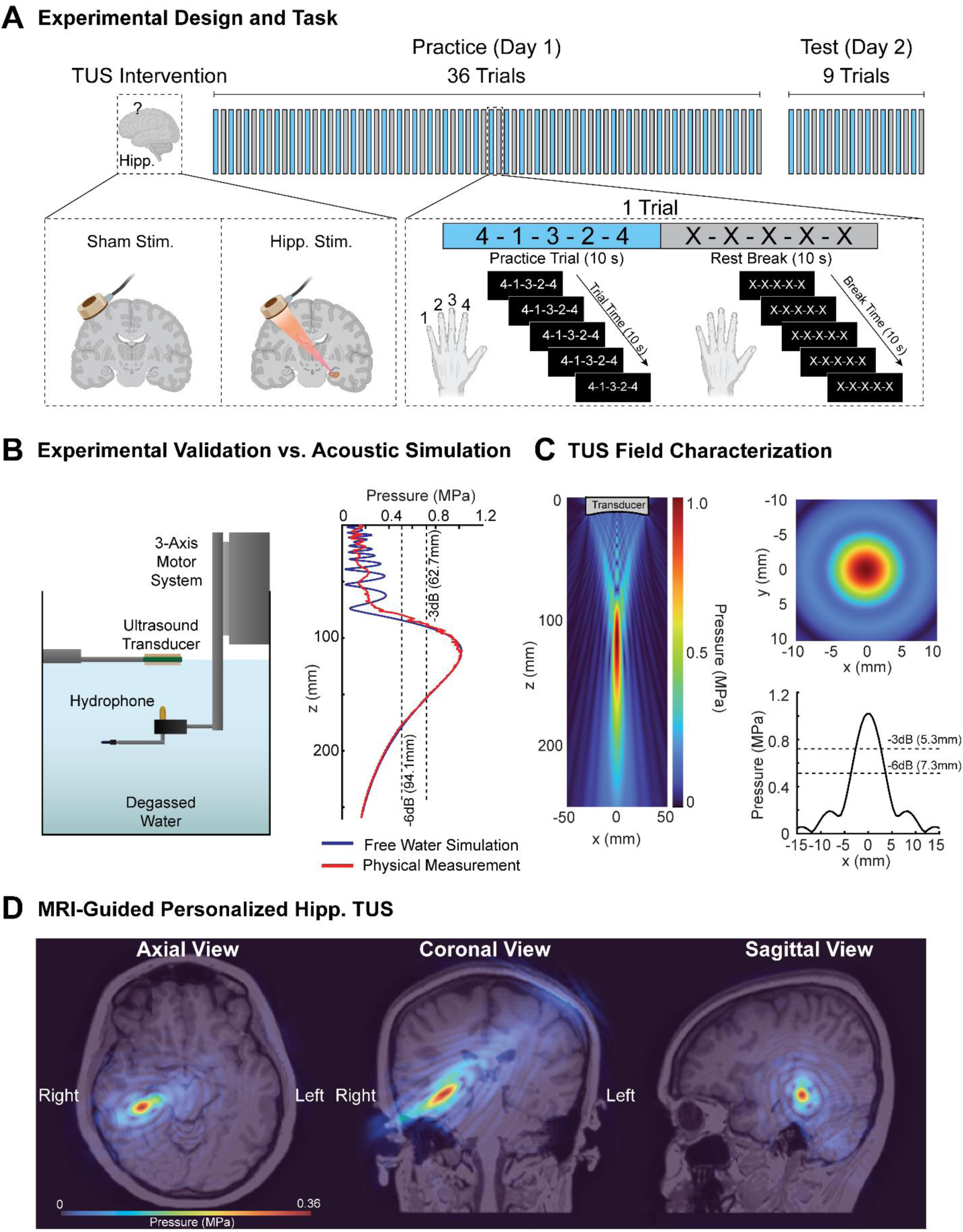
Experimental Design, Acoustic Characterization, and Personalized Hippocampal TUS Targeting. (**A**) Experimental design and motor sequence task. Participants were assigned to hippocampal TUS or sham stimulation. On Day 1, stimulation was followed by 36 trials of motor sequence learning. Approximately 24 h later, participants returned for a 9-trial Day 2 retest without additional stimulation. Each trial consisted of a 10-s practice period, during which participants repeatedly typed the five-element sequence (4-1-3-2-4), followed by a 10-s rest period. (**B**) Hydrophone-based validation of the acoustic simulation. Left, schematic of the hydrophone measurement setup, with the transducer and hydrophone positioned in degassed water using a three-axis motorized system. Right, axial pressure profiles obtained from physical hydrophone measurements and free-water acoustic simulation, showing close agreement in focal location and pressure distribution. (**C**) Characterization of the simulated acoustic field. Left, two-dimensional pressure distribution along the axial direction of the ultrasound beam. Top right, pressure distribution at the focal plane. Bottom right, lateral pressure profile through the focus, with the −3-dB full-width at half-maximum (FWHM) and −6-dB widths indicated. (**D**) Representative individualized MRI-guided hippocampal TUS targeting. Simulated intracranial pressure fields are overlaid on the participant’s anatomical MRI in axial, coronal, and sagittal views, showing the simulated acoustic field centered on the intended posterior hippocampal target.

### Acoustic Field Characterization and Personalized TUS Targeting

To validate our acoustic simulation, we first characterized the acoustic output of the 0.5-MHz transducer using hydrophone measurements in a degassed water tank (Figure 1B). The physical measurements and simulation were closely aligned, as demonstrated by the overlap between the measured (red) and simulated (blue) axial pressure profiles (Figure 1B). The measured focal depth was 111.4 mm, closely matching the simulated target depth of 112.0 mm, which corresponded to the average depth required to target the posterior hippocampus. At a target spatial-peak pulse-average intensity (I_sppa_) of 35 W/cm², the measured peak pressure was 1.03 MPa, compared with the simulated value of 1.02 MPa. This agreement supported the use of the acoustic simulation model to characterize the three-dimensional pressure field (Figure 1C). The simulation showed a collimated acoustic field with an elongated, ellipsoidal focal spot. The axial full-width at half-maximum (FWHM) was 62.7 mm, with a -6 dB width of 94.1 mm. In contrast, the focus was more confined in the lateral plane, with a FWHM of just 5.3 mm and a -6 dB width of 7.3 mm (Figure 1C). Together, these measurements and simulations show that the protocol generated an acoustic field centered at the target depth, with substantially greater confinement in the lateral than axial dimension.

To ensure safe and accurate delivery of TUS in vivo, we employed an MRI-guided personalized TUS targeting pipeline for each participant (Figure 1D). This process utilized individual high-resolution T1-weighted anatomical images that were converted into subject-specific pseudo-CTs to model skull density and morphology, allowing transcranial acoustic simulations to guide transducer placement and assess whether the acoustic focus reached the posterior hippocampus (Figure 1D). Across all participants, the mean transducer focal depth was 112.0 ± 6.3 mm with an I_sppa_ of 35.9 ± 7.7 W/cm². After accounting for skull attenuation, the simulations predicted a mean in situ Isppa at the posterior hippocampus of 4.3 ± 0.06 W/cm² and a corresponding pressure of 0.36 ± 0.003 MPa. The simulations also indicated that the stimulation parameters remained within established mechanical and thermal safety limits. The predicted mechanical index (MI) was 0.52 ± 0.03, well below the FDA diagnostic-ultrasound output limit, and the predicted thermal dose (CEM43) was negligible (3.88 × 10⁻⁴ ± 2.7 × 10⁻⁶), indicating a minimal risk of tissue heating (Aubry et al., 2025). This individualized procedure allowed us to account for differences in skull anatomy across participants and to consistently center the acoustic focus on the intended target region.

### TUS Protocol Produces Sustained Suppression of Motor Cortex Excitability

To first validate the neuromodulatory effect of our TUS protocol, we examined its effects on primary motor cortex (M1) excitability (Figure 2A). We measured motor-evoked potentials (MEPs) elicited by single-pulse TMS at baseline and again at 5 and 30 minutes following either active TUS or sham stimulation over M1. A 2 × 3 mixed-design analysis of variance (ANOVA), with Group (Active, Sham) as the between-subjects factor and Time (Baseline, 5 min, 30 min) as the within-subjects factor, revealed a significant Group × Time interaction (F(2, 76) = 4.22, p = 0.018), indicating that changes in M1 excitability over time differed between active and sham stimulation. Planned paired-samples t-tests within the active condition showed a sustained suppression of MEP amplitude (Figure 2B). Specifically, MEPs were significantly reduced from a baseline of 1.15 ± 0.30 mV to 0.92 ± 0.30 mV at 5 minutes (t(19) = 3.26, p = 0.004) and remained significantly suppressed at 0.96 ± 0.29 mV at 30 minutes (t(19) = 2.86, p = 0.010). MEP amplitudes did not differ between 5 and 30 minutes (p = 0.482). In contrast, MEP amplitudes following sham stimulation did not significantly change from baseline (1.16 ± 0.58 mV) at either 5 minutes (1.22 ± 0.65 mV; t(19) = −0.66, p = 0.516) or 30 minutes (1.24 ± 0.67 mV; t(19) = −1.20, p = 0.247) (Figure 2C). These results show that the TUS parameters used in this study can produce a sustained reduction in cortical excitability that persists for at least 30 minutes, encompassing the duration of the main behavioral experiment.

**Figure 2.**
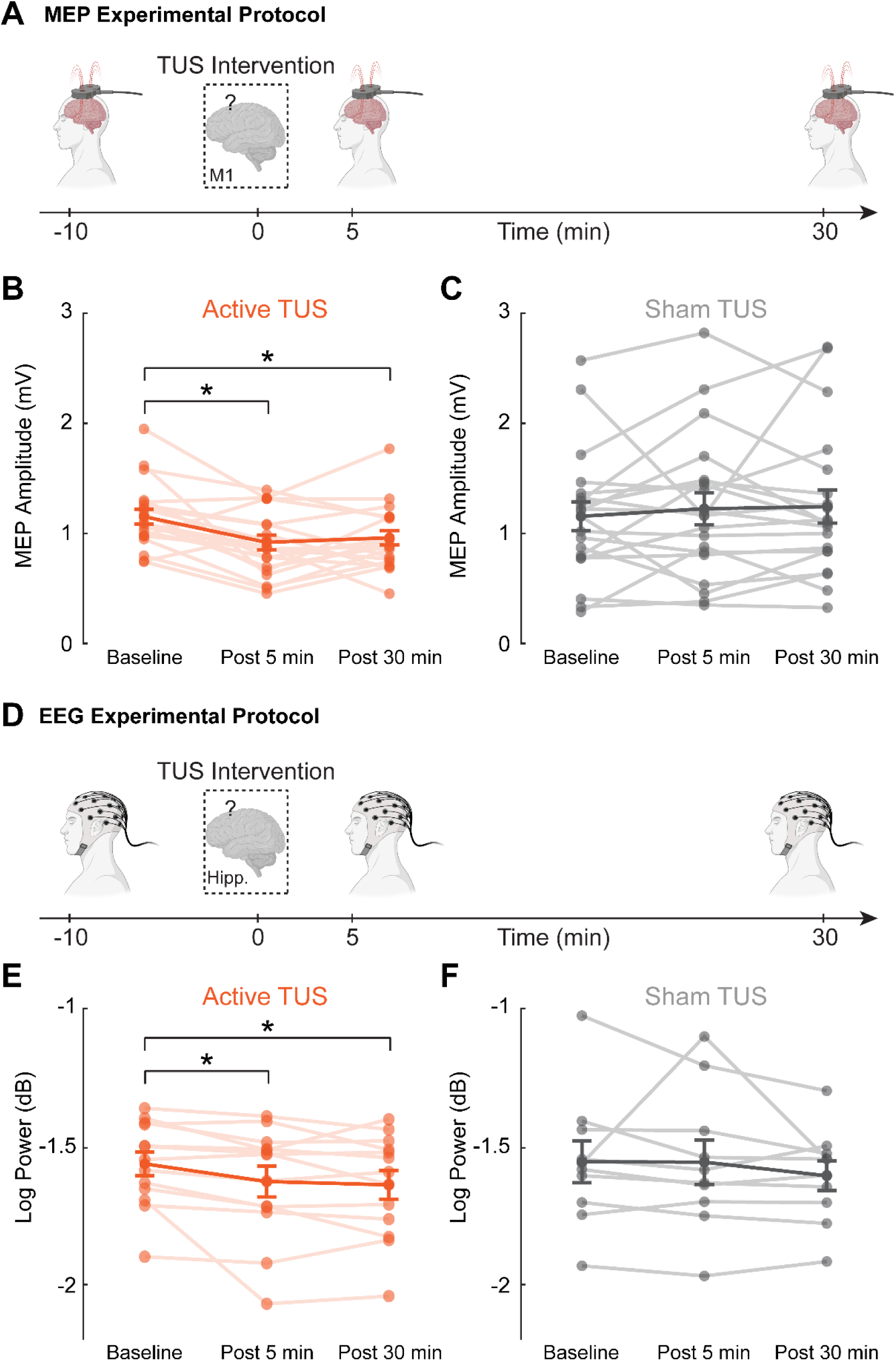
Physiological Effects of TUS Assessed by MEPs and EEG. (**A**) Experimental protocol for the M1 validation experiment. MEPs were recorded at Baseline and approximately 5 and 30 min after active or sham TUS over M1. (**B**) MEP amplitudes in the active TUS group (*N* = 20). Individual participant values are shown across Baseline, Post5, and Post30, with group means superimposed. MEP amplitude was reduced at both Post5 and Post30 relative to Baseline. (**C**) MEP amplitudes in the sham group (*N* = 20). No significant changes were observed across the three time points. (**D**) Experimental protocol for the EEG validation experiment. Resting-state EEG was recorded at Baseline and approximately 5 and 30 min after active or sham TUS targeting the posterior hippocampus. (**E**) Frontocentral gamma-band power in the active hippocampal TUS group (*N* = 13). Gamma power decreased at Post5 and Post30 relative to Baseline. (**F**) Frontocentral gamma-band power in the sham group (*N* = 10). No significant changes from Baseline were observed. Individual participant values and group means are shown. * indicates *p* < 0.05 for planned within-group comparisons.

### Frontocentral Gamma Activity Decreases Following Hippocampal TUS

Having established the suppressive effect of the TUS protocol over M1, we next targeted the posterior hippocampus and examined frontocentral gamma-band activity (Figure 2D). Based on the inhibitory effect observed over M1 and the known interactions between the hippocampus and frontal cortical regions, we hypothesized a priori that active hippocampal TUS would reduce frontocentral gamma-band activity. A 2 × 3 mixed-design ANOVA with Group (Active, Sham) as the between-subjects factor and Time (Baseline, 5 min, 30 min) as the within-subjects factor showed a significant main effect of Time (F(2, 42) = 3.55, p = 0.038), but no significant Group × Time interaction (F(2, 42) = 0.81, p = 0.454). Planned comparisons showed that frontocentral gamma power decreased following active TUS (Figure 2E), from −1.56 ± 0.15 dB at baseline to −1.63 ± 0.20 dB at 5 minutes (t(12) = 2.30, p = 0.040), and remained lower at 30 minutes (−1.64 ± 0.19 dB; t(12) = 2.65, p = 0.021). Gamma power did not differ between the 5- and 30-minute time points (t(12) = 0.60, p = 0.558). In the sham condition, gamma-band power remained stable from baseline (−1.55 ± 0.24 dB) at both 5 minutes (−1.56 ± 0.26 dB; t(9) = 0.03, p = 0.976) and 30 minutes (−1.61 ± 0.17 dB; t(9) = 1.64, p = 0.136) (Figure 2F). Together, these results indicate that frontocentral gamma activity decreased following active hippocampal TUS and remained reduced for at least 30 minutes.

### Hippocampal TUS Shifts Learning from Offline to Online Gains

To determine how hippocampal TUS altered the learning process, we separated Day 1 performance gains into micro-online and micro-offline components, following previous work on rapid motor memory consolidation (Bönstrup et al., 2019). Micro-online gains reflected changes in tapping speed during each 10-s practice period, whereas micro-offline gains reflected changes that emerged across the intervening rest periods (Figure 3A,D). A 2 × 2 mixed-design ANOVA with Group (Hippocampal TUS, Sham) as the between-subjects factor and Gain Type (micro-online, micro-offline) as the within-subjects factor revealed a significant Group × Gain Type interaction (*F*(1, 38) = 18.07, *p* < 0.001), indicating that hippocampal TUS altered the distribution of performance gains between active practice and rest.

**Figure 3.**
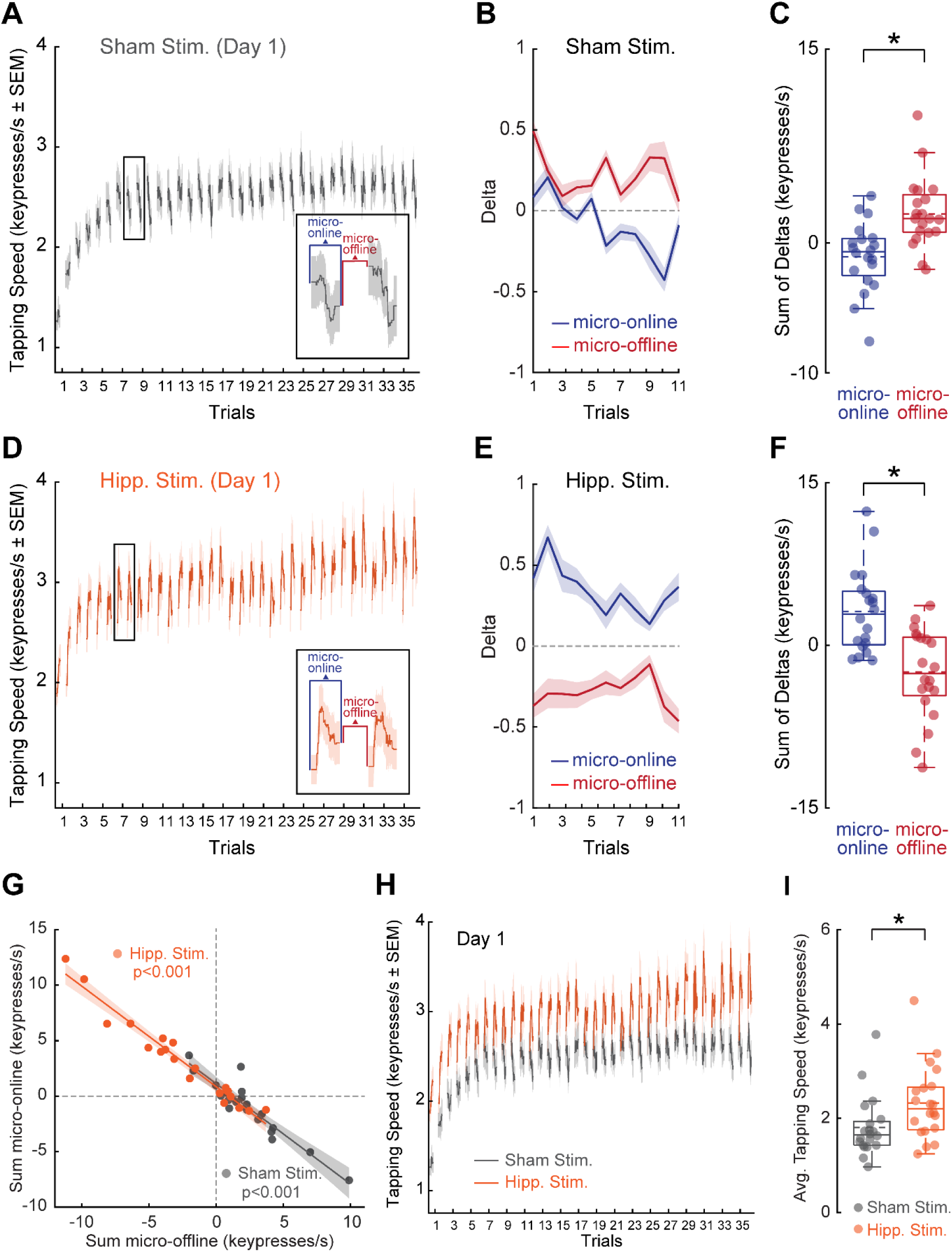
Hippocampal TUS Shifts Day 1 Learning from Offline to Online Gains and Increases Overall Performance. (**A**) Tapping speed (keypresses/s, mean ± SEM) across the 36 Day 1 learning trials in the sham group. The inset illustrates the calculation of micro-online changes during active practice and micro-offline changes across the intervening rest period. (**B**) Micro-online and micro-offline performance changes across the first 11 Day 1 trials in the sham group. (**C**) Summed micro-online and micro-offline gains across the first 11 trials in the sham group. Micro-offline gains were greater than micro-online gains. (**D**) Tapping speed across the 36 Day 1 learning trials in the hippocampal TUS group. (**E**) Micro-online and micro-offline performance changes across the first 11 Day 1 trials following hippocampal TUS. (**F**) Summed micro-online and micro-offline gains across the first 11 trials in the hippocampal TUS group. Micro-online gains were greater than micro-offline gains. (**G**) Relationship between summed micro-online and micro-offline gains in the sham and hippocampal TUS groups. Strong inverse correlations were observed in both groups, while hippocampal TUS shifted the distribution toward greater online and lower offline gains. (**H**) Group-averaged Day 1 tapping performance for the sham and hippocampal TUS groups. (**I**) Mean Day 1 tapping performance. Participants receiving hippocampal TUS showed higher overall tapping speed than those receiving sham stimulation. Shaded areas indicate ± SEM. Box plots show individual participant values, medians, interquartile ranges, and whiskers extending to non-outlier values. * indicates *p* < 0.05.

In the sham group, summed micro-offline gains were positive (2.35 ± 2.74 keypresses/s), whereas micro-online gains were slightly negative (−1.01 ± 2.66 keypresses/s; Figure 3B,C). Micro-offline gains were significantly greater than micro-online gains (t(19) = 2.83, p = 0.011), consistent with the typical pattern in which early performance improvements emerge predominantly during brief rest periods. Hippocampal TUS produced the opposite pattern (Figure 3E,F). Micro-online gains were positive (3.15 ± 3.83 keypresses/s), whereas micro-offline gains were negative (−2.54 ± 4.13 keypresses/s), with micro-online gains significantly greater than micro-offline gains (*t*(19) = 3.22, *p* = 0.004). Direct comparisons between groups showed that hippocampal TUS produced greater micro-online gains (t(38) = 3.99, p < 0.001) and lower micro-offline gains (t(38) = −4.42, p < 0.001) than sham stimulation. Together, these results indicate that hippocampal TUS shifted the expression of early learning from gains occurring during rest toward gains occurring during active practice.

Despite these different learning patterns, micro-online and micro-offline gains were inversely correlated within both groups (Figure 3G). This relationship was observed in the sham group (r = −0.93, R² = 0.87, p < 0.001) and in the hippocampal TUS group (r = −0.98, R² = 0.95, p < 0.001). Participants receiving sham stimulation tended to occupy the higher-offline/lower-online portion of this relationship, whereas those receiving hippocampal TUS were shifted toward higher online and lower offline gains. These results suggest that online and offline gains are tightly coupled during early skill learning, with hippocampal TUS shifting how performance improvements are expressed between rest and active practice.

### Hippocampal TUS Enhances Overall Task Performance

We next asked whether this redistribution of learning was accompanied by higher overall task performance. Despite the reduction in micro-offline gains, participants receiving hippocampal TUS showed higher tapping performance across Day 1 practice than those receiving sham stimulation (Figure 3H,I). Average tapping speed was 3.08 ± 0.78 keypresses/s following hippocampal TUS, compared with 2.56 ± 0.64 keypresses/s following sham stimulation (t(38) = 2.31, p = 0.026). This higher tapping speed was not accompanied by reduced accuracy. Accuracy was similarly high in the hippocampal TUS (97.59 ± 2.24%) and sham groups (97.85 ± 1.13%; *t*(38) = −0.47, *p* = 0.644). Thus, the shift toward gains expressed during active practice was accompanied by higher overall performance during initial learning without a reduction in accuracy.

### Independent Replication Confirms the Shift from Offline to Online Learning

In a separate replication experiment, we included a baseline practice period before stimulation to account for potential differences in motor performance between groups (Figure 4A). Participants first completed 24 randomized sequence trials matched in structure to the learning sequence, allowing us to assess baseline tapping performance without sequence-specific learning. They then received either hippocampal TUS or sham stimulation and completed the same 36-trial motor sequence learning task used in the initial experiment. Baseline tapping performance was similar between groups (Figure 4B–D), averaging 1.65 ± 0.42 keypresses/s in the hippocampal TUS group and 1.63 ± 0.40 keypresses/s in the sham group (*t*(18) = 0.09, *p* = 0.930). Thus, the two groups began the sequence-learning task with comparable motor performance. Baseline accuracy was also comparable between groups (hippocampal TUS: 95.67 ± 2.88%; sham: 94.17 ± 3.45%; *t*(18) = 1.06, *p* = 0.305). Accuracy remained comparable during subsequent sequence learning (hippocampal TUS: 97.65 ± 1.64%; sham: 96.49 ± 2.14%; *t*(18) = 1.36, *p* = 0.190).

**Figure 4.**
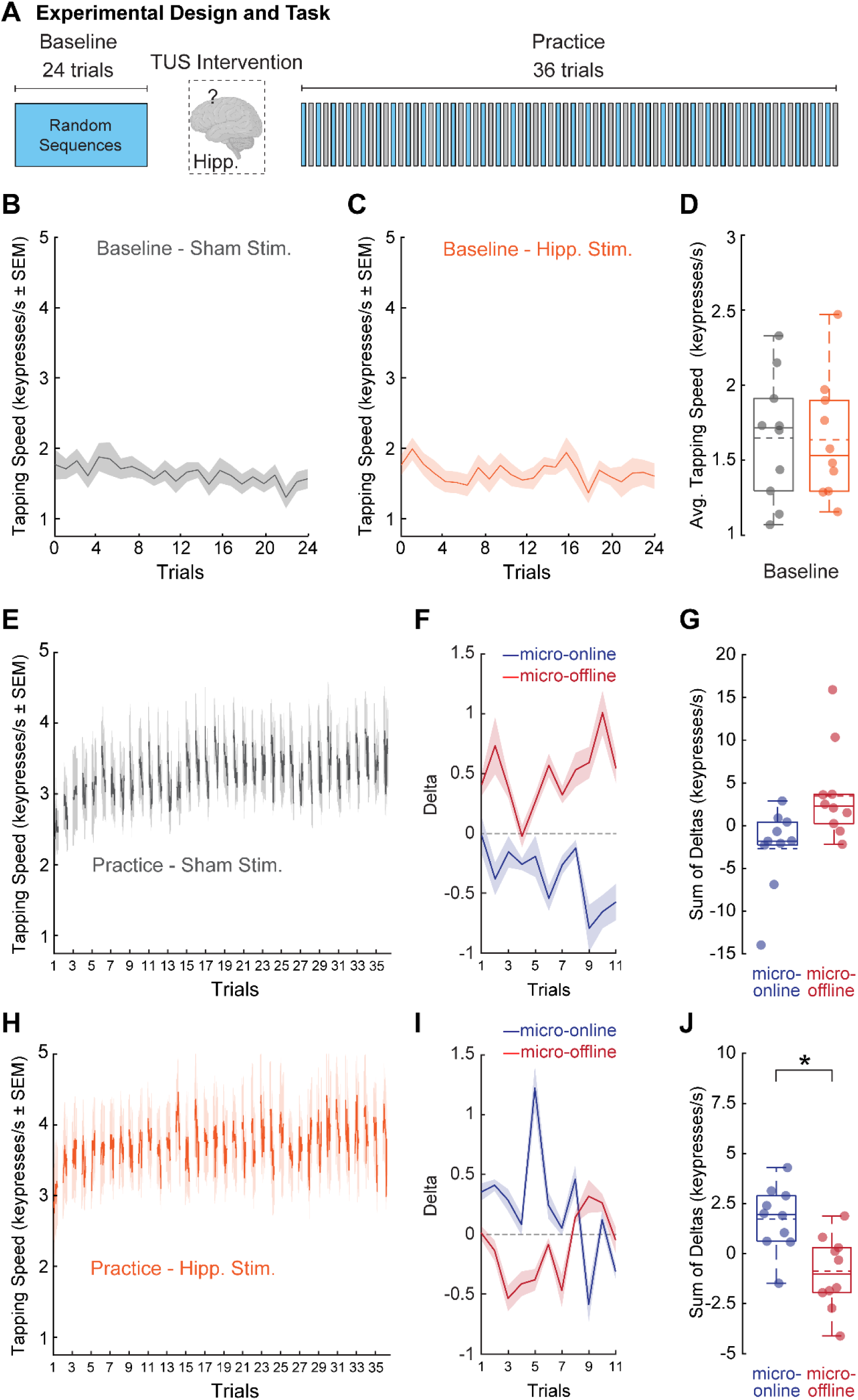
Independent Replication of the Shift from Offline to Online Learning. (**A**) Design of the independent replication experiment. Participants first completed 24 random-sequence trials to assess baseline motor performance before receiving hippocampal TUS or sham stimulation. Stimulation was followed by 36 trials of motor sequence learning. (**B, C**) Baseline tapping performance across the 24 random-sequence trials in the sham (**B**) and hippocampal TUS (**C**) groups. (**D**) Mean baseline tapping performance did not differ between groups, indicating comparable motor performance before stimulation and sequence learning. (**E**) Tapping speed across the 36 sequence-learning trials in the sham group. (**F**) Micro-online and micro-offline performance changes across the first 11 sequence-learning trials in the sham group. (**G**) Summed micro-online and micro-offline gains across the first 11 trials in the sham group. The difference between gain types did not reach statistical significance. (**H**) Tapping speed across the 36 sequence-learning trials in the hippocampal TUS group. (**I**) Micro-online and micro-offline performance changes across the first 11 sequence-learning trials following hippocampal TUS. (**J**) Summed micro-online and micro-offline gains across the first 11 trials in the hippocampal TUS group. Micro-online gains were significantly greater than micro-offline gains, reproducing the shift observed in the main experiment. Shaded areas indicate ± SEM. Box plots show individual participant values, medians, interquartile ranges, and whiskers extending to non-outlier values. * indicates *p* < 0.05.

We then tested whether the shift between micro-online and micro-offline learning observed in the initial experiment was reproduced in this cohort. A 2 × 2 mixed-design ANOVA with Group (Hippocampal TUS, Sham) as the between-subjects factor and Gain Type (micro-online, micro-offline) as the within-subjects factor revealed a significant Group × Gain Type interaction (F(1, 18) = 6.96, p = 0.017). In the sham group, micro-offline gains were positive (3.72 ± 5.45 keypresses/s), whereas micro-online gains were negative (−2.52 ± 4.77 keypresses/s; Figure 4E–G), although the difference between the two gain types was not significant (t(9) = 1.94, p = 0.084). In contrast, participants receiving hippocampal TUS showed positive micro-online gains (1.74 ± 1.63 keypresses/s) and negative micro-offline gains (−0.95 ± 1.82 keypresses/s; Figure 4H–J), with micro-online gains significantly greater than micro-offline gains (t(9) = 2.54, p = 0.032). Direct comparisons between groups showed greater micro-online gains following hippocampal TUS than sham stimulation (*t*(18) = 2.67, *p* = 0.015), together with lower micro-offline gains (*t*(18) = −2.57, *p* = 0.019). Together, these results reproduced the shift from offline to online learning observed in the initial experiment, with both groups starting from comparable baseline motor performance.

### Altered Learning Dynamics and Higher Performance Persist 24 Hours Later

We next asked whether the altered learning pattern observed on Day 1 persisted when participants returned approximately 24 hours later. A 2 × 2 mixed-design ANOVA with Group (Hippocampal TUS, Sham) as the between-subjects factor and Gain Type (micro-online, micro-offline) as the within-subjects factor revealed a significant Group × Gain Type interaction (*F*(1, 38) = 15.59, *p* < 0.001), indicating that the distribution of learning between practice and rest continued to differ between groups on Day 2.

In the sham group, micro-online gains were positive (1.13 ± 2.21 keypresses/s), whereas micro-offline gains were slightly negative (−0.42 ± 2.05 keypresses/s; Figure 5A–C), although the difference between the two gain types was not significant (*t*(19) = 1.65, *p* = 0.115). In contrast, participants who had received hippocampal TUS continued to show a clear online-dominant pattern (Figure 5D–F), with positive micro-online gains (5.27 ± 4.21 keypresses/s) and negative micro-offline gains (−4.12 ± 3.70 keypresses/s). Micro-online gains were significantly greater than micro-offline gains in this group (*t*(19) = 5.35, *p* < 0.001). Direct comparisons between groups showed greater micro-online gains following hippocampal TUS (*t*(38) = 3.89, *p* < 0.001) and lower micro-offline gains (*t*(38) = −3.92, *p* < 0.001).

**Figure 5.**
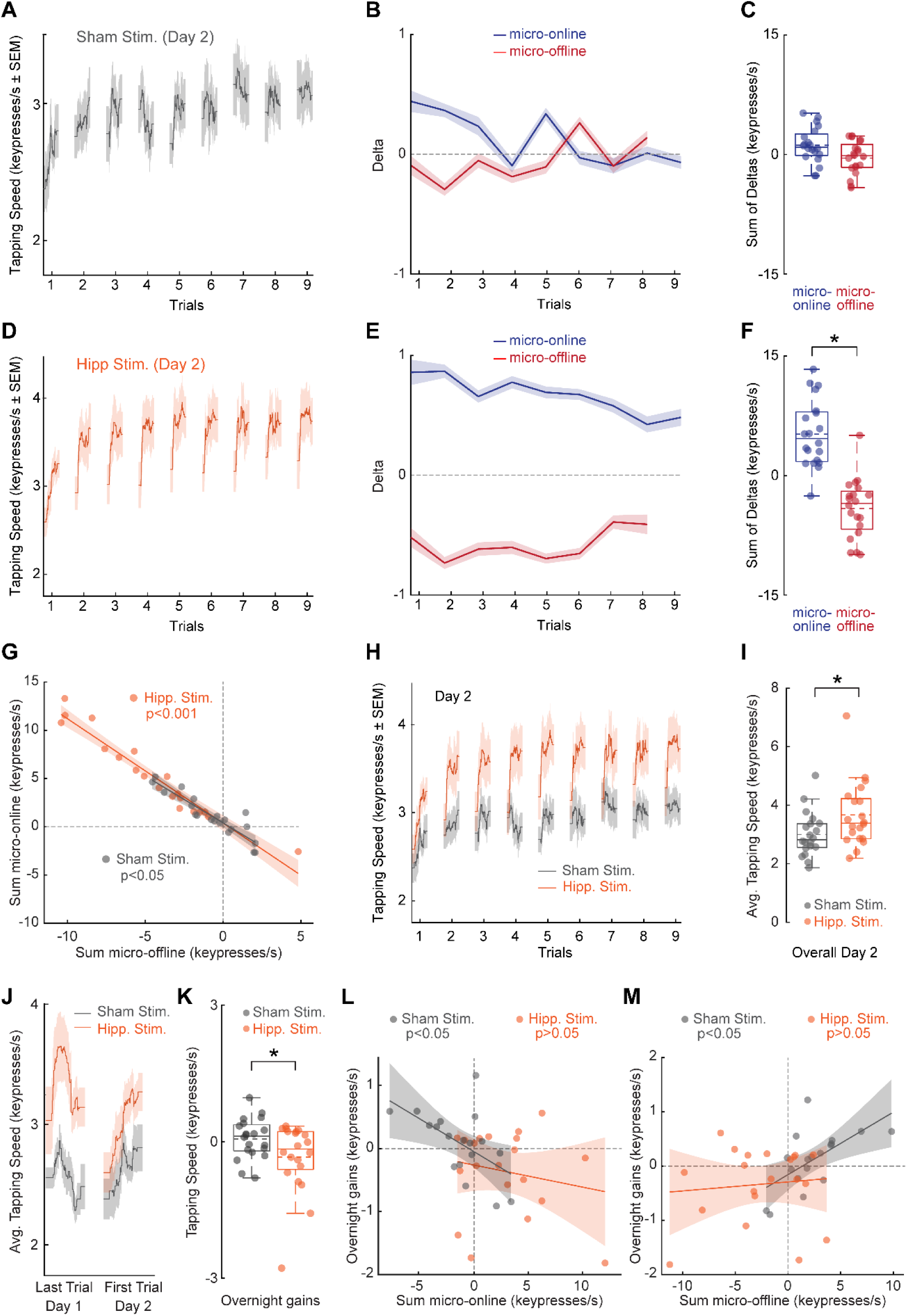
Altered Learning Dynamics and Higher Performance Remain Evident 24 Hours Later. (**A**) Tapping speed (keypresses/s, mean ± SEM) across the nine Day 2 trials in the sham group. (**B**) Micro-online and micro-offline performance changes across Day 2 in the sham group. (**C**) Summed micro-online and micro-offline gains in the sham group. The difference between gain types was not significant. (**D**) Tapping speed across the nine Day 2 trials in the hippocampal TUS group. (**E**) Micro-online and micro-offline performance changes across Day 2 following hippocampal TUS. (**F**) Summed micro-online and micro-offline gains in the hippocampal TUS group. Micro-online gains were significantly greater than micro-offline gains. (**G**) Relationship between summed micro-online and micro-offline gains on Day 2. Strong inverse correlations were observed in both the sham and hippocampal TUS groups. (**H**) Group-averaged tapping performance across the Day 2 session. (**I**) Mean Day 2 tapping performance. The hippocampal TUS group maintained higher overall performance than the sham group. (**J**) Change in performance across the overnight interval, shown from the final trial of Day 1 to the first trial of Day 2. (**K**) Overnight change in tapping performance. Overnight change did not differ from zero within either group, but the change differed significantly between the hippocampal TUS and sham groups. (**L**) Relationship between Day 1 micro-online gains and overnight change. A significant negative correlation was observed in the sham group, whereas no significant relationship was observed following hippocampal TUS. (**M**) Relationship between Day 1 micro-offline gains and overnight change. A significant positive correlation was observed in the sham group, whereas no significant relationship was observed following hippocampal TUS. Shaded regions in correlation plots indicate 95% confidence intervals. Shaded regions in performance curves indicate ± SEM. Box plots show individual participant values, medians, interquartile ranges, and whiskers extending to non-outlier values. * indicates *p* < 0.05.

The inverse relationship between micro-online and micro-offline gains was also evident on Day 2 (Figure 5G). The two measures were negatively correlated in both the sham group (*r* = −0.92, R² = 0.84, *p* < 0.001) and the hippocampal TUS group (*r* = −0.97, R² = 0.94, *p* < 0.001), suggesting that the coupling between gains during practice and rest remained present 24 hours later. Importantly, the altered learning dynamics were accompanied by higher overall performance on Day 2 (Figure 5H,I). Participants who had received hippocampal TUS had an average tapping speed of 3.63 ± 1.11 keypresses/s, compared with 2.99 ± 0.75 keypresses/s in the sham group (*t*(38) = 2.14, *p* = 0.039). Accuracy remained high and did not differ between the hippocampal TUS (98.53 ± 1.42%) and sham groups (98.34 ± 1.73%; *t*(38) = 0.38, *p* = 0.707). Thus, the higher performance observed during initial learning was still present 24 hours later, together with the shift toward gains expressed during active practice.

### Day 1 Learning Dynamics Predict Overnight Change in the Sham Group

We next examined whether the pattern of learning on Day 1 was related to changes in performance across the overnight interval. Overnight change was quantified from performance at the end of Day 1 to the beginning of Day 2 (Figure 5J,K). The sham group showed little average change overnight (0.06 ± 0.44 keypresses/s; *t*(19) = 0.65, *p* = 0.523), whereas the hippocampal TUS group showed a modest decrease (−0.34 ± 0.77 keypresses/s; *t*(19) = −1.98, *p* = 0.063). The overnight change differed between groups (*t*(38) = −2.04, *p* = 0.049).

Within the sham group, the learning pattern on Day 1 was related to subsequent overnight change. Greater micro-online gains were associated with less overnight improvement (r = −0.54, R² = 0.29, p = 0.014; Figure 5L), whereas greater micro-offline gains were associated with greater overnight improvement (r = 0.61, R² = 0.37, p = 0.005; Figure 5M). These relationships were not evident in participants who received hippocampal TUS, for either micro-online gains (r = −0.19, R² = 0.04, p = 0.410) or micro-offline gains (r = 0.10, R² = 0.01, p = 0.681). Together, these findings suggest that hippocampal TUS altered the relationship between Day 1 learning dynamics and overnight consolidation.

## Discussion

This study provides causal evidence that the human hippocampus contributes to the balance between online and offline processes during early motor learning. By applying non-invasive TUS to the posterior hippocampus, we changed the temporal pattern of skill acquisition. Specifically, hippocampal TUS reduced micro-offline gains during rest while increasing performance gains during active practice. We replicated this shift from offline to online learning in an independent cohort with comparable baseline motor performance. Importantly, this redistribution of learning did not impair overall skill acquisition. Instead, hippocampal TUS enhanced task performance during learning, and this higher performance remained evident 24 hours later. Together, these results suggest that the hippocampus helps determine how learning is distributed between practice and rest and that this balance can be altered through targeted neuromodulation.

Converging methodological and physiological evidence supports successful targeting of the intended posterior hippocampal region and a physiological effect of stimulation. First, hydrophone measurements closely matched the simulated acoustic field, supporting the accuracy of our computational model. We then used individualized, MRI-based acoustic simulations to account for differences in skull anatomy across participants and guide placement of the acoustic focus within the posterior hippocampus. Physiologically, applying the suppressive stimulation protocol over M1 produced a sustained reduction in corticospinal excitability for at least 30 minutes, providing a physiological benchmark for the direction and duration of the stimulation effect (Bao et al., 2024). When the same TUS parameters were applied to the posterior hippocampus, frontocentral gamma-band activity decreased and remained lower for up to 30 minutes. Although scalp EEG cannot directly measure local hippocampal activity, the combination of acoustic validation, individualized targeting, and physiological changes following stimulation provides converging support for successful modulation of the intended hippocampal target.

The most striking behavioral finding was the redistribution of learning from micro-offline to micro-online gains. Consistent with previous work, our sham group reproduced the well-established “micro-consolidation” pattern (Bönstrup et al., 2019), in which a substantial portion of early performance improvements emerges during brief rest periods. This process has been linked to the reactivation of learning-related neural patterns, with neuroimaging studies showing that waking hippocampo-neocortical replay and hippocampal activity during rest predict subsequent behavioral gains (Jacobacci et al., 2020; Buch et al., 2021). Consistent with this, prior work involving patients with hippocampal damage or network disruption via prefrontal stimulation has shown that interfering with hippocampal function attenuates these micro-offline gains (Gann et al., 2023; Mylonas et al., 2024). Our findings extend this work by demonstrating that modulating hippocampal activity not only reduces rest-related gains but also shifts performance improvements toward active practice. Following hippocampal TUS, micro-online gains became positive, while micro-offline gains inverted to become negative on average. This pattern suggests that the hippocampus may help regulate how learning is distributed between practice and rest, rather than exclusively supporting offline consolidation.

The independent replication strengthened this finding by addressing potential baseline differences between groups. Because the initial experiment used a between-subjects design, inherent differences in motor ability before stimulation could have influenced the learning trajectories. In the replication cohort, participants completed a random-sequence baseline period before stimulation, allowing us to assess motor performance without sequence-specific learning. Baseline tapping speed and accuracy were comparable between groups. Participants receiving hippocampal TUS again showed a shift toward positive micro-online and negative micro-offline gains. Thus, the same change in learning dynamics was reproduced in participants who began with comparable baseline motor performance, supporting the reliability of the main behavioral finding.

One possible explanation for this redistribution comes from theories of competitive memory systems, which describe interactions between hippocampal and striatal learning processes (Poldrack and Packard, 2003). Reducing reliance on one system can, under some conditions, increase reliance on another. For instance, hippocampal lesions can enhance performance on sequence-learning tasks in rodents (Eckart et al., 2012), and individuals with hippocampal damage can show preserved or enhanced implicit learning (Reber and Squire, 1998). Similarly, placing the declarative memory system under cognitive load can bias learning toward striatal mechanisms in healthy adults (Foerde et al., 2006). Corticostriatal networks are also well established as contributors to motor sequence acquisition (Doyon et al., 2003; Doyon and Benali, 2005; Dayan and Cohen, 2011). In this context, reducing hippocampal influence may have allowed greater engagement of practice-dependent corticostriatal processes. Our study did not measure striatal activity directly, however, so this remains a hypothesis that will require direct testing.

Micro-online and micro-offline gains were inversely correlated on both Day 1 and Day 2. Participants in the sham group tended toward higher offline and lower online gains, whereas hippocampal TUS shifted this pattern toward higher online and lower offline gains. Rather than eliminating the relationship between the two components, TUS appeared to shift where participants fell along it. This suggests that hippocampal modulation changes how performance improvements are distributed between practice and rest. Importantly, this shift was accompanied by higher overall performance without a reduction in accuracy. Participants receiving hippocampal TUS performed better across Day 1, despite showing markedly reduced micro-offline gains, and this higher level of performance remained evident 24 hours later. Thus, reducing rest-related gains did not compromise skill acquisition or its persistence. Instead, in this task, greater expression of learning during active practice was associated with higher overall performance. This finding is consistent with the broader view that motor memory depends on multiple interacting neural systems whose contributions change across learning and consolidation (Doyon et al., 2003; Dayan and Cohen, 2011; Krakauer et al., 2019).

The overnight results further distinguish these two learning patterns. In the sham group, greater micro-offline gains on Day 1 were associated with greater overnight improvement, whereas greater micro-online gains were associated with less overnight improvement. This relationship is consistent with the idea that processes expressed during brief waking rest may be related to subsequent stabilization across longer intervals (Walker et al., 2002; Doyon et al., 2009). Neither relationship was evident following hippocampal TUS. Hippocampal TUS also did not enhance overnight consolidation itself. Although performance showed a modest decline across the overnight interval, the TUS group remained at a higher performance level on Day 2, reflecting the higher performance achieved during initial learning. Thus, the persistent behavioral effect appears to reflect the higher level of skill acquired on Day 1 rather than enhanced overnight consolidation. Together, these findings suggest that hippocampal TUS altered the relationship between Day 1 learning dynamics and overnight consolidation. This pattern is consistent with models in which multiple learning systems contribute to the acquisition and stabilization of motor skills (White and McDonald, 2002; Poldrack and Packard, 2003).

The interpretation of micro-offline gains as a direct measure of memory consolidation remains debated. Recent behavioral work has suggested that these gains may partly reflect recovery from fatigue, motor preparation, or other transient performance factors (Das et al., 2025). At the same time, recent evidence argues against a purely fatigue- or planning-based account of early skill-learning dynamics. Iwane et al. (2026) showed that within-trial performance patterns changed systematically with learning and were related to hippocampal theta–gamma coupling, providing additional evidence that hippocampal memory processes contribute to early motor skill acquisition. Our findings do not resolve this debate completely, but they add causal evidence for a hippocampal contribution. Targeted hippocampal TUS altered the balance between micro-online and micro-offline gains, and this effect was reproduced in an independent cohort with comparable baseline motor performance. This finding is consistent with previous work linking micro-offline gains to hippocampal activity during rest and waking hippocampo-neocortical replay (Jacobacci et al., 2020; Buch et al., 2021). Taken together, micro-offline gains are likely multifactorial, reflecting both transient performance effects and neural processes related to memory stabilization.

The ability to shift learning toward active practice may have translational relevance. In neurorehabilitation, where fatigue can limit the amount of effective practice (Hinkle et al., 2017), increasing gains expressed during active training could improve training efficiency. Similar questions may be relevant to populations with impaired sleep-dependent memory processes, including older adults with age-related changes in memory consolidation (Mander et al., 2017). Several limitations should also be considered. First, the present experiments involved healthy adults performing a relatively simple sequence task, and it remains unclear whether the effect generalizes to more complex skills or clinical populations. Second, although EEG showed a physiological response following hippocampal TUS, scalp recordings do not directly measure local hippocampal activity. Because we did not record neural activity during the motor-learning task itself, the proposed involvement of corticostriatal processing remains inferential. Third, an active control stimulation site would provide stronger evidence for anatomical specificity. Although individualized acoustic modeling and image-guided targeting allowed the acoustic focus to be centered within the posterior hippocampus, they cannot establish that neuromodulatory effects were restricted to this region. Given the elongated axial extent of the acoustic field, neighboring medial temporal structures may also have received acoustic energy and contributed to the observed effects. Fourth, the subject-specific acoustic simulations relied on T1-derived pseudo-CT images rather than acquired CT. Although this approach avoids ionizing radiation and enables individualized targeting, uncertainty in estimated skull properties may introduce error in the modeled estimates of transcranial pressure and attenuation. Finally, the replication cohort was smaller than the initial cohort. Future studies should test the effect in larger and more diverse samples and examine longer-term retention, interference, and transfer.

In conclusion, our findings identify the human hippocampus as an important contributor to how motor learning is distributed between active practice and brief rest. Hippocampal TUS shifted learning toward online gains, and this effect was reproduced in an independent cohort with comparable baseline performance. This altered learning pattern was accompanied by higher overall performance that remained evident 24 hours later. Together, these results show that the balance between online and offline motor learning is flexible and can be altered through targeted modulation of the hippocampus.

## Materials and Methods

### Participants

A total of 72 healthy volunteers participated in the study. The original cohort consisted of 52 participants, and an additional 20 participants were recruited for the independent replication experiment. Within the original cohort, 40 participants completed the motor cortex TUS validation experiment (active TUS, N = 20; sham, N = 20), 23 completed the EEG validation experiment following hippocampal TUS (active TUS, N = 13; sham, N = 10), and 40 completed the main behavioral experiment (hippocampal TUS, N = 20; sham, N = 20). The independent replication cohort consisted of 20 new participants who were randomly assigned to hippocampal TUS (*N* = 10) or sham stimulation (*N* = 10). All participants were free of medications known to alter brain excitability and reported no history of neurological or psychiatric disorders. Hand dominance was determined by self-report. For participants who completed more than one experimental session, a minimum interval of five days was maintained between sessions to minimize potential carry-over effects. The study protocol was approved by the local Institutional Review Board (IRB), and all participants provided written informed consent before participation. All procedures were conducted in accordance with the Declaration of Helsinki.

### Experimental Design

The study used randomized, between-subject designs to examine the physiological and behavioral effects of TUS. The study consisted of two validation experiments and two behavioral experiments. The validation experiments were designed to characterize the physiological effects of the stimulation protocol. In the motor cortex validation experiment, 40 participants were assigned to active TUS (N = 20) or sham stimulation (N = 20). Corticospinal excitability was assessed before stimulation and again approximately 5 and 30 min after stimulation. In a separate EEG validation experiment, 23 participants received either active hippocampal TUS (N = 13) or sham stimulation (N = 10), and EEG activity was assessed before stimulation and at approximately 5 and 30 min after stimulation. The main behavioral experiment included 40 participants randomly assigned to hippocampal TUS (N = 20) or sham stimulation (N = 20). The experiment was conducted across two consecutive days (Figure 1A). On Day 1, participants received active or sham stimulation immediately before completing 36 trials of a motor sequence learning task. Approximately 24 h later, participants returned for Day 2 and completed a 9-trial retest without additional stimulation. This design allowed us to examine both the immediate effects of hippocampal TUS on learning during practice and rest and whether differences in performance remained evident the following day. An independent replication experiment was conducted in 20 new participants, who were randomly assigned to hippocampal TUS (N = 10) or sham stimulation (N = 10). To assess baseline motor performance independently of sequence-specific learning, participants first completed 24 trials of random-sequence practice before stimulation. They then received active or sham hippocampal TUS and completed the same 36-trial motor sequence learning task used in the main experiment (Figure 4A). The replication experiment was completed in a single session and did not include a Day 2 retest. The baseline period was included to determine whether the two groups showed comparable motor performance before stimulation and before learning of the repeated sequence.

### Motor Sequence Task

Participants performed a procedural motor sequence task adapted from previous work (Bönstrup et al., 2019). Using the non-dominant hand, they repeatedly typed the five-element sequence 4-1-3-2-4 as quickly and accurately as possible on a standard QWERTY keyboard (4 = index, 3 = middle, 2 = ring, 1 = little finger). The sequence remained visible during practice, and an asterisk appeared after each keypress to provide immediate visual feedback. Each trial consisted of a 10-s practice period followed by a 10-s rest period (Figure 1A). During practice, participants repeated the sequence for the full 10 s, with the number of completed repetitions determined by individual performance speed. During rest, the sequence was replaced by five “X” symbols (X-X-X-X-X), and participants were instructed to keep their hand still and fixate on the screen. In the main behavioral experiment, participants completed 36 learning trials on Day 1 and 9 retest trials approximately 24 h later on Day 2, with no additional TUS administered on Day 2. Testing was conducted at approximately the same time of day across sessions (within ±2 h) to reduce potential circadian effects. In the independent replication experiment, participants first completed 24 randomized sequence trials matched in structure to the learning sequence, allowing us to assess baseline tapping performance without sequence-specific learning. They then received active or sham hippocampal TUS and completed the same 36-trial learning task used on Day 1 of the main experiment.

### Transcranial Ultrasound Stimulation (TUS)

For hippocampal stimulation, TUS was delivered using a four-element, 500-kHz spherical ultrasound transducer (DPX-500-4CH-008; 64.0-mm diameter, 63.2-mm radius of curvature; Sonic Concepts). For the M1 validation experiment, a 500-kHz CTX transducer (CTX-500-4CH-082; Sonic Concepts) was used because its focal geometry was better suited to the shallower cortical target. Apart from the transducer used to accommodate the different target depths, all sonication parameters were identical between the M1 validation and hippocampal stimulation protocols. The stimulation protocol consisted of 20-ms ultrasound bursts delivered at 5 Hz for 80 s (400 bursts total), corresponding to a 10% duty cycle. Individualized acoustic simulations were used to determine the transducer placement and stimulation parameters required to target the posterior hippocampus while accounting for interindividual differences in skull anatomy. Acoustic and thermal exposure estimates were maintained within established safety recommendations (MI ≤ 1.9, I_sppa_ ≤ 190 W/cm², and CEM43 ≤ 0.25; FDA, 2019; Aubry et al., 2025; Murphy et al., 2025). Participant-level estimates of in situ I_sppa_, peak pressure, mechanical index (MI), and thermal dose (CEM43) were obtained from the acoustic simulations and are summarized in the Results.

### Acoustic Measurements Using a Calibrated Hydrophone

The acoustic output of the transducer was independently characterized using a calibrated hydrophone scanning system (AIMS III; Onda Corp.) in a degassed water tank at room temperature (Figure 1B). An HGL-0200 hydrophone (Onda Corp.) was mounted on a three-axis motorized positioning system, and the hydrophone signal was passed through a preamplifier to an oscilloscope synchronized with the transducer output. A three-dimensional scan was first performed to locate the point of maximum acoustic pressure and thereby identify the geometric focus. Following localization of the focus, a high-resolution one-dimensional scan was performed along the axial (z) direction to characterize the beam profile. Peak pressure and spatial characteristics of the focal field were quantified from these measurements and compared with the corresponding simulated acoustic field.

### Anatomical MRI Acquisition

Prior to TUS, a high-resolution T1-weighted structural MRI was acquired for each participant using a magnetization-prepared rapid gradient-echo (MPRAGE) sequence on a 3-T Siemens Prisma scanner. Images were acquired at 1-mm isotropic resolution (matrix = 256 × 256 × 192; repetition time [TR] = 2300 ms; echo time [TE] = 2.28 ms; flip angle = 8°). The anatomical images were used to generate individualized pseudo-CT images for acoustic modeling and to guide neuronavigated placement of the TUS transducer using Brainsight (Rogue Research, Canada).

### Pseudo-CT Generation for Acoustic Modeling

Because skull morphology and composition substantially affect transcranial ultrasound propagation through attenuation, reflection, and refraction (Pinton et al., 2011), individualized skull models were generated for acoustic simulation. Rather than acquiring CT scans, which involve ionizing radiation, pseudo-CT images were generated from each participant’s T1-weighted MRI using an open-source deep-learning toolbox (Yaakub et al., 2023). The method uses a pretrained convolutional neural network to estimate skull geometry and density from structural MRI. Prior to pseudo-CT generation, T1-weighted images were bias-field corrected and preprocessed according to the requirements of the model. The resulting pseudo-CT images were then used as the anatomical input for individualized acoustic and thermal simulations.

### TUS Simulation

Individualized acoustic simulations were performed using k-Plan (Brainbox Ltd., UK) to account for the effects of skull anatomy on ultrasound propagation. The simulations were based on the pseudo-CT images generated from each participant’s structural MRI. Pseudo-CT Hounsfield units were converted to material properties for acoustic modeling using a lower threshold of 300 HU and an upper limit of 2000 HU. The simulation volume was segmented into skull bone (>1,150 kg/m³), soft tissue (1,045 kg/m³), and water (996 kg/m³). For each participant, the transducer position and orientation were iteratively optimized so that the simulated acoustic focus was centered within the posterior hippocampal target defined on the individual anatomical MRI. Simulations were performed in the time domain, and pressure amplitude and phase were calculated after the acoustic field reached steady state. The resulting pressure fields were used to estimate focal location, in situ pressure and intensity, mechanical index, and thermal exposure and to guide the subsequent neuronavigated transducer placement.

### Personalized TUS Targeting

The posterior hippocampus was selected a priori as the stimulation target. Participants performed the sequence-learning task with the non-dominant hand, and the posterior hippocampus contralateral to the task hand was selected as the stimulation target based on its proposed involvement in visuospatial and sequence-related learning processes. Individualized acoustic simulations were used to determine the optimal transducer position and orientation for each participant. During stimulation, the transducer was coupled to the scalp using ultrasound gel and tracked with the Brainsight neuronavigation system after registration of the participant’s head to the anatomical MRI. The transducer position and orientation were adjusted to match the configuration identified by the individualized acoustic simulation. Once the planned targeting configuration was achieved, the 80-s TUS protocol was delivered. Transducer position was monitored throughout stimulation to minimize displacement from the planned trajectory. For sham stimulation, the positioning, coupling, neuronavigation, and experimental procedures were identical to the active condition, but ultrasound output was not delivered. To reduce the possibility that participants could distinguish active from sham stimulation based on auditory cues, continuous Gaussian white noise was presented throughout the stimulation period in both conditions.

### Motor-Evoked Potential Recording

To characterize the effect of TUS on cortical excitability, MEPs were elicited using single-pulse TMS before and after stimulation of M1. TMS was delivered using a DuoMAG MP-Dual system (Brainbox Ltd.) with a 70-mm figure-of-eight coil. The coil was positioned tangentially over the M1 hand area, with the handle directed posteriorly and laterally at approximately 45° to the midline. The stimulation site was identified individually and monitored throughout the experiment using neuronavigation co-registered to each participant’s anatomical MRI. TMS intensity was set to the minimum intensity required to evoke MEPs with a peak-to-peak amplitude greater than 1 mV in at least 5 of 10 consecutive trials and was maintained at the same intensity throughout the experiment (Bao and Lei, 2025; Bao et al., 2024). MEPs were recorded from the contralateral first dorsal interosseous (FDI) muscle using disposable Ag-AgCl surface electrodes in a belly-tendon montage. EMG signals were amplified and band-pass filtered from 30 to 2000 Hz using a Neurolog System (Digitimer) and digitized at 10 kHz using a CED Micro 1401. MEPs were collected at three time points: before TUS (Baseline), approximately 5 min after stimulation (Post5), and approximately 30 min after stimulation (Post30) (Figure 2A). At each time point, 24 single-pulse TMS trials were recorded.

### EEG Recording

Resting-state EEG was recorded using an actiCHamp amplifier and actiCAP active electrode system (Brain Products GmbH, Germany) with BrainVision Recorder software. A modified 64-channel electrode configuration was used to accommodate placement of the TUS transducer. Twelve electrodes (F7, F3, FC5, M1, T7, C3, M2, FC3, C5, PO6, FT7, and PO7) were omitted, leaving 52 available EEG channels. Participants were seated comfortably and instructed to remain relaxed, minimize head, facial, and body movements, and maintain visual fixation throughout the recordings. Electrode impedance was checked before each recording. EEG was collected at three time points: before stimulation (Baseline), approximately 5 min after stimulation (Post5), and approximately 30 min after stimulation (Post30). At each time point, two 2-min resting-state recordings were obtained.

### Behavioral Data Analysis

Behavioral data were processed using custom scripts in MATLAB (The MathWorks). The primary behavioral measure was tapping speed, expressed in keypresses per second. For each correctly executed sequence, tapping speed was calculated from the time intervals between adjacent keypresses. Trial performance was defined as the mean tapping speed across all correctly executed sequences within each 10-s practice period. Response accuracy was calculated for each participant as the percentage of sequences completed correctly within each experimental phase. For the main behavioral experiment, accuracy was calculated separately for Day 1 learning and Day 2 retest. For the replication experiment, accuracy was calculated separately for the pre-stimulation baseline and post-stimulation sequence-learning periods. Within-session learning was separated into micro-online and micro-offline components following previous work (Bönstrup et al., 2019). Micro-online gain was calculated as the tapping speed of the last correct sequence minus that of the first correct sequence within each practice period, with positive values indicating improvement during active practice. Micro-offline gain was calculated as the tapping speed of the first correct sequence of the subsequent trial minus that of the last correct sequence of the preceding trial, with positive values indicating improvement across the intervening rest period. Consistent with Bönstrup et al. (2019), analyses of micro-online and micro-offline learning on Day 1 focused on the first 11 trials, corresponding to the early phase of skill acquisition. The same approach was used for the post-stimulation learning period in the independent replication experiment. For the Day 2 retest, all nine trials were included. Micro-online and micro-offline gains were summed separately across the relevant trials to provide participant-level measures of learning expressed during practice and rest. Overall task performance was calculated as the mean tapping speed across all trials within a session. For the independent replication experiment, baseline motor performance was calculated from the 24 random-sequence trials completed before stimulation. For the main behavioral experiment, overnight change was calculated as the tapping speed of the first trial on Day 2 minus that of the final trial on Day 1, with positive values indicating overnight improvement and negative values indicating a decline.

### Neurophysiological Data Analysis

MEP data were processed using Signal software (Cambridge Electronic Design). Peak-to-peak MEP amplitude was calculated for each response and averaged across the 24 trials collected at Baseline, Post5, and Post30 to obtain a mean MEP amplitude for each participant at each time point. EEG data were processed using EEGLAB and MNE. Resting-state recordings were analyzed separately at Baseline, Post5, and Post30. Non-EEG channels and channels identified as flat or disconnected were excluded. Signals were notch filtered at 60 Hz to reduce line noise, band-pass filtered from 1 to 100 Hz, resampled to 512 Hz, and re-referenced to the common average reference. Gamma-band power (30–50 Hz) was estimated for each channel using Welch’s power spectral density method and log10-transformed. Spectral estimates were calculated separately for the two recordings at each time point and then averaged. The primary EEG analysis focused on the midline frontocentral electrodes Fz, FCz, and Cz. Gamma-band power was averaged across these electrodes to generate a single measure of frontocentral gamma activity for each participant at each time point.

### Statistical Analysis

Statistical analyses were performed using SPSS Statistics (IBM Corp.). For the M1 validation experiment, MEP amplitudes were analyzed using a 2 × 3 mixed-design ANOVA with Group (active TUS, sham) as the between-subjects factor and Time (Baseline, Post5, Post30) as the within-subject factor. Planned paired-samples *t*-tests examined changes across time within each group. For the EEG validation experiment, frontocentral gamma-band power was analyzed using a 2 × 3 mixed-design ANOVA with Group (hippocampal TUS, sham) as the between-subjects factor and Time (Baseline, Post5, Post30) as the within-subject factor. Planned paired-samples *t*-tests examined changes from Baseline to Post5 and Post30, as well as differences between Post5 and Post30, within each group. For the main behavioral experiment, micro-online and micro-offline gains were analyzed separately for Day 1 and Day 2 using 2 × 2 mixed-design ANOVAs with Group (hippocampal TUS, sham) as the between-subjects factor and Gain Type (micro-online, micro-offline) as the within-subject factor. Follow-up paired-samples *t*-tests compared micro-online and micro-offline gains within each group, and independent-samples *t*-tests compared the two groups for each gain type. Overall task performance and accuracy were compared between groups using independent-samples *t*-tests. Associations between micro-online and micro-offline gains were assessed separately within each group using Pearson’s correlation coefficient (*r*). For the independent replication experiment, baseline tapping speed and accuracy were compared between groups using independent-samples *t*-tests. Accuracy during subsequent sequence learning was also compared between groups using an independent-samples *t*-test. Learning during the subsequent sequence task was analyzed using the same 2 × 2 Group × Gain Type mixed-design ANOVA and follow-up comparisons used for Day 1 of the main experiment. Overnight change in the main behavioral experiment was assessed using one-sample *t*-tests against zero within each group and an independent-samples *t*-test between groups. Pearson correlations were used to examine the relationships between Day 1 micro-online and micro-offline gains and overnight change separately within each group.

